# Grapevine leaf size influences vine canopy temperature

**DOI:** 10.1101/2022.07.07.499216

**Authors:** Zoë Migicovsky, Joel F. Swift, Zachary Helget, Laura L. Klein, Anh Ly, Matthew Maimaitiyiming, Karoline Woodhouse, Anne Fennell, Misha Kwasniewski, Allison J. Miller, Daniel H. Chitwood, Peter Cousins

**Affiliations:** Department of Plant, Food, and Environmental Sciences, Faculty of Agriculture, Dalhousie University, Truro, Nova Scotia, B2N 5E3, Canada; Department of Biology, Acadia University, Wolfville, Nova Scotia, B4P 2R6, Canada; Department of Biology, Saint Louis University, St. Louis, MO, 63103, USA; Agronomy, Horticulture, and Plant Science, South Dakota State University, Brookings, SD, 57007, USA; Department of Natural and Applied Sciences, Missouri State University, Springfield, MO, 65897, USA; Division of Food Sciences, University of Missouri, Columbia, MO, 65211, USA; Department of Food Sciences, The Pennsylvania State University, University Park, PA, 16802, USA; Department of Horticulture, Michigan State University, East Lansing, MI 48823 USA; Department of Computational Mathematics, Science & Engineering, Michigan State University, East Lansing, MI, 48823, USA; E. & J. Gallo Winery, Modesto, CA 95354, USA

**Keywords:** ampelography, grapevine, leaf morphology, leaf temperature, leaf shape, *Vitis*

## Abstract

**Premise:** Grapevine leaves have diverse shapes and sizes. Their shape and size is known to be influenced by many factors including genetics, vine phytosanitary status, environment, leaf and vine age, and node position on the shoot. In order to determine the importance of grapevine leaf shape and size to canopy temperature, we examined the relationship in five seedling populations grown in a vineyard in California, USA.

**Methods:** All of the populations had one parent with compound leaves of the *Vitis piasezkii* type and each population had a different second parent with non-compound leaves. In previous work, we measured leaf shape and size using 21 homologous landmarks. Here, we paired these morphology data with measurements taken using an infrared thermometer to measure the temperature of the canopy. By recording time of sampling and canopy temperature, we were able to determine which vines were cooler or hotter than expected, using a linear model.

**Results:** We established a relationship between leaf size and canopy temperature: vines with larger leaves were cooler than expected. In contrast, leaf shape was not strongly correlated with variation in temperature.

**Conclusions:** Ultimately, these findings indicate that vines with larger leaves may contribute to the reduction of overall vine canopy temperature, but further work is needed to determine if this is due to variation in leaf size, differences in the openness of the canopy, or other related traits.

## Introduction

Grapevine (*Vitis* spp.) leaves have diverse shapes and sizes. The field of ampelography (“vine” + “writing”) is dedicated to the study of grapevine leaves, enabling the identification of both species and individual cultivars (Galet, 1979; Chitwood et al., 2016a; Chitwood, 2021). The shape and size of grapevine leaves is influenced by genetics (Chitwood et al., 2014; Demmings et al., 2019), vine phytosanitary status (Klein et al., 2017), environment (Chitwood et al., 2016b, 2021), leaf age as well as node position (Chitwood et al., 2016a; b; Bryson et al., 2020), rootstock (Migicovsky et al., 2019; Harris et al., 2021), and many other factors.

As the primary photosynthetic organs of the plant, increasing leaf size increases photosynthetic potential of the plant. However, the increase in leaf size may also have negative consequences: larger leaves with a thicker boundary layer may slow heat loss, increasing respiration at a rate greater than the increase in photosynthesis (Givnish, 1987; Westoby et al., 2002). Both the size of an individual leaf and the combined size of all leaves, also known as the total leaf area, may have an effect on plant growth and health. For example, in grapevine, the higher water demand for transpiration associated with a larger total leaf area may increase vine water stress, and as a result, reduce yield (Mirás-Avalos et al., 2017).

In addition, the overall temperature of a vine or canopy temperature is an important consideration. Canopy temperature can be measured by infrared thermometry, including remote thermal imaging, and is therefore non-invasive and non-destructive (Leinonen and Jones, 2004; Giménez-Gallego et al., 2021). Temperature can influence many developmental processes in grapevines, with higher temperatures accelerating development, including the timing of budbreak, bloom, and onset of fruit ripening, a particular concern in the face of climate change (Keller and Tarara, 2010; Parker et al., 2011). While ambient temperature plays a critical role, canopy temperature is also important. For example, it is the bud temperature, rather than air temperature, that determines the timing of budbreak (Keller and Tarara, 2010).

Canopy temperature can not only influence vine development, but also performance. In a controlled study of three different canopy temperatures for *Vitis vinifera* ‘Semillon’ vines, differences in up to 3°C in mean canopy temperature over the growing season impacted the vines. In particular, reproductive growth was impacted with berry expansion and sugar accumulation being the highest at the lowest temperature, although yield was generally not affected (Greer and Weedon, 2019).

The role of canopy temperature is not restricted to vine development and characteristics, it can also be an indicator of water availability. Transpiration contributes to cooling of the leaf and can be particularly important in sunlit leaves. When the air is still or wind speeds are very low, the amount of transpiration occurring has a larger effect on leaf temperature. In cases where leaf temperature approaches a lethal temperature, the effect of transpiration on leaf temperature can be critical to survival (Gates, 1964).

Many grapevines are irrigated and efficient management of water stress in both vines and other plants requires that the grower knows when water stress has begun and how much water to apply. Canopy temperature can be used to assess plant water status using the crop water stress index (CWSI) which is calculated based on the difference between canopy and air temperature (Cohen et al., 2005). For example, work in common bean (*Phaseolus vulgaris* L.) using the CWSI found that the above ground measurements of air and leaf temperature were as effective at predicting plant water stress as soil-based measurements (Durigon and de Jong van Lier, 2013).

Similar to other crops, the CWSI may also be used in grapevines to determine the need for and effect of irrigation (Ahi et al., 2015). Thus, canopy temperature both plays an important role in vine development and is a critical indicator of vine water status. How the shape and size of grapevine leaves interact with the environment to influence canopy temperature is poorly understood. Given that grapevine leaf shape is at least partly controlled by genetics (Chitwood et al., 2014; Demmings et al., 2019), if particular leaf shapes or sizes have a positive impact on canopy temperature, such as keeping vine temperature low in hotter climates and reducing the need for irrigation, this could be a desirable target for grape breeders.

In this study, we examined the importance of grapevine leaf shape and size to canopy temperature. Using an infrared thermometer to measure the temperature of the canopy we determined which vines were cooler or hotter than expected. We established a relationship between leaf size and leaf temperature: vines with larger leaves were cooler than expected. In contrast, leaf shape was not strongly correlated with variation in vine temperature. These findings provide evidence that leaf size, but not shape, may contribute to canopy temperature.

## Materials and Methods

### Experimental design

Leaves were sampled from seedlings of five biparental *Vitis* populations located in San Joaquin Valley, Madera County, California. As described in Migicovsky et al. (In Press), and copied here for convenience, the populations consisted of a total of 500 seedlings. 450 seedlings had DVIT 2876 as one parent. The remaining 50 seedlings had DVIT 2876 as a grandparent. DVIT 2876 ‘Olmo b55-19’ is a compound-leafed accession from the USDA-ARS National Clonal Germplasm repository, suspected to include *Vitis piasezkii* Maximowicz, as one of its parents (or grandparents). Thus, all of the populations had one parent with compound leaves of the *V. piasezkii* type and each population had a different second parent with non-compound leaves. The populations were created to examine variation in leaf lobing and the resulting progeny from each cross had a range of leaf shapes from very lobed to entire.

Sampled populations (Figure 1A) included 125 individuals from a DVIT 2876 x unnamed *Vitis vinifera* selection cross (Pop1), 100 individuals from a DVIT 2876 x a different unnamed *Vitis vinifera* selection cross (Pop2), 150 individual from a DVIT 2876 x unnamed *Vitis* hybrid cross (Pop3), 75 individual from a DVIT 2876 x a different unnamed *Vitis* hybrid cross (Pop4), and 50 individuals from a seedling (DVIT 2876 x unnamed *Vitis vinifera* selection) x DVIT 3374 (*Vitis mustangensis* Buckley) cross (Pop5). Selections used in these crosses are unnamed because they are the result of breeding crosses. Vines were planted in 2017. They were trained to a unilateral cordon and spur pruned.

**Figure 1.**
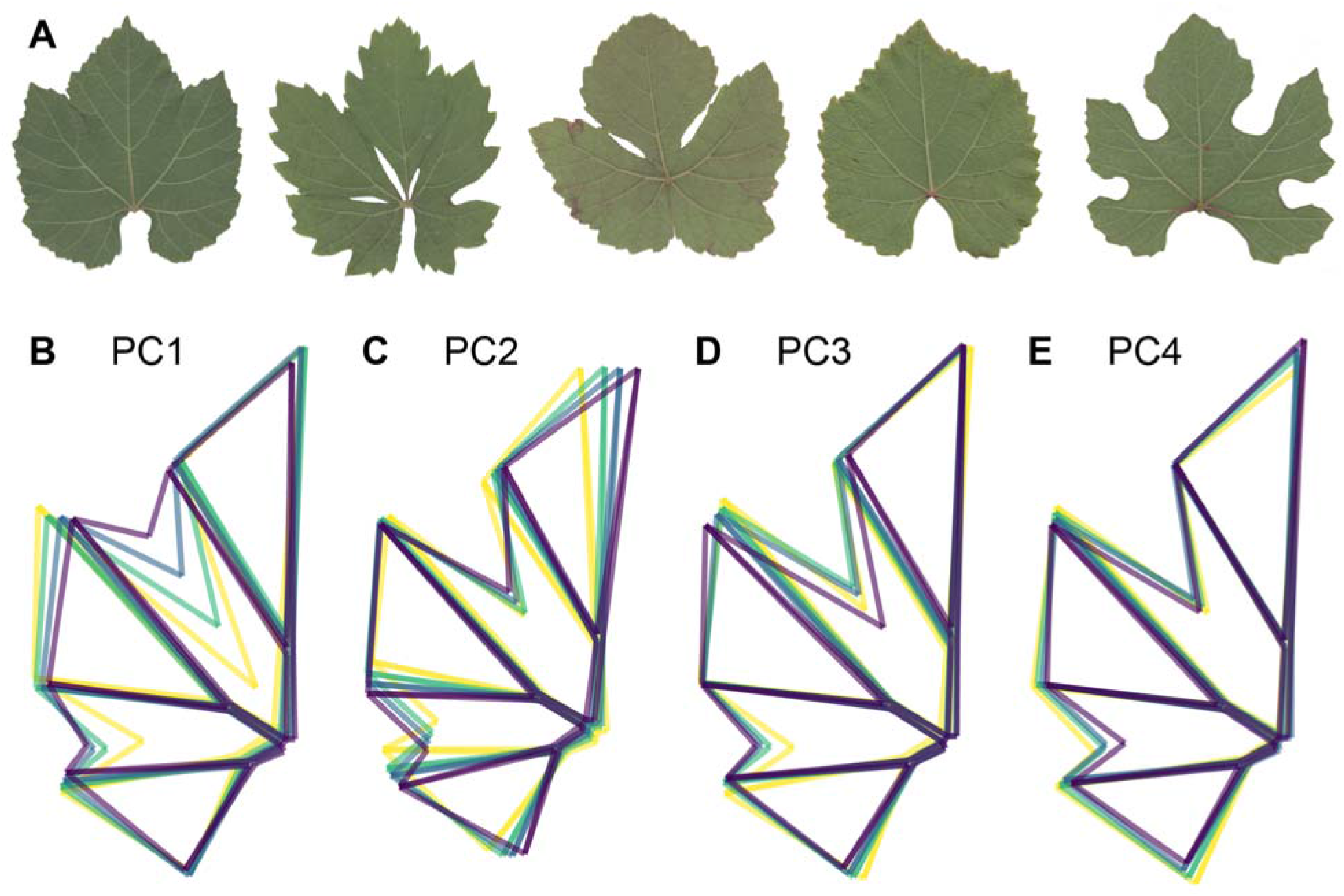
Real and eigenleaves showing variation in shape across the populations sampled. (A) A leaf from each of the five populations (Pop1 to 5 shown from left to right) showing the range of lobing present across the accessions. Given the extensive range of lobing within a population, the leaf shown is not representative of a particular population, but rather used to show the range across all populations. (B-E) For each PC quartile, a mean leaf is plotted, with the lowest PC quartile shown in yellow, increasing in color to dark blue across quartiles. PC1 to PC4, explaining a total of 78% of the variance in leaf shape, are shown.

### Sampling

Three representative leaves were collected and scanned from each vine across June and July in 2018, and then again across June and July 2019. For full details of leaf collection and scanning, see Migicovsky et al. (In Press).

Leaf temperature measurements took place twice in 2018 (July 19 and August 10) and twice in 2019 (July 24 and August 1). For three of the four dates, measurements were taken from approximately 9 AM to 11 AM, but on one date (July 24 2019) measurements were taken from approximately 11:30 AM to 1:30 PM. Measurements were taken using an infrared thermometer (Extech 42515 InfraRed Thermometer) to measure the temperature of the canopy. These measurements were taken by using the thermometer to scan across the outside of the canopy and then recording the mean temperature of the vine.

In most cases, the exact time of the measurement was also recorded. However, in some cases, time was recorded every few vines. In these instances, the time of sampling was interpolated by dividing the difference in time between two measurements by the number of measurements taken between those two times, and adding that to the initial time. For example, if a vine was measured at 9:11 AM and a second vine was measured at 9:13 AM, the unlabelled vine in between those two measurements would have been recorded as 9:12 AM. If breaks were needed or a new row of measurements began, the time was always recorded on the last and first vine measured before/after those periods.

Vines which were too small to accurately measure, for example, those with only a few leaves, or those that were dead, were not measured. For the purposes of this study, vines were reduced to only those with at least one leaf scanned for shape in both 2018 and 2019, and with canopy temperature measurements recorded at all four timepoints. As a result, the total number of unique accessions across all time points used for the analyses in this study was 388 out of the 500 vines initially planted.

Weather data for each of the dates was downloaded from an on-site weather station, which included temperature measurements taken once per hour for a total of 24 measurements per day. Precipitation was also recorded although there was no precipitation during the sampling days.

### Data analysis

Image analysis of the leaf scans is fully described in Migicovsky et al. (In Press) and scans are available on Dryad (Migicovsky et al., 2022). Briefly, leaves were analyzed using 21 landmarks as previously described by (Chitwood et al., 2016b, 2021; Bryson et al., 2020). Leaf area was calculated using the shoelace algorithm, which calculates the area of a polygon using the landmarks as vertices, following previously described methods (Chitwood et al., 2021). In addition, we calculated the ratio of vein to blade area of each leaf as well as the degree of distal and proximal lobing. Following adjustment using a generalized Procrustes analysis in the shapes package in R (Dryden, 2021) principal components analysis (PCA) was performed to determine the primary sources of variation in leaf shape.

Subsequent analyses were performed in R and code is available at the following GitHub repository https://github.com/zoemigicovsky/grape_leaf_temp. All visualizations were performed using ggplot2 v3.3.5 (Wickham, 2016).

For this study, instead of using shape measurements for individual leaves, morphometric values were averaged across measurements taken from a vine in a given year in order to be able to connect average leaf shape and size with canopy temperature measurements. Given that representative leaves were selected, even vines with fewer than three leaves sampled were retained and averaged (when more than one leaf was sampled).

Only three measurements exceeded 105 °F, and only one value was less than 64 °F, and thus, these were considered likely errors and removed from the dataset. Temperature measurements were converted from Fahrenheit to Celsius using the weathermetrics version 1.2.2 package in R (Anderson and Peng, 2012) for downstream analyses.

Using the broom package in R (Robinson et al., 2021), a linear model was performed for each date, to determine the effect of time of sampling on canopy temperature (Equation 1):

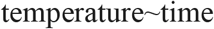

by extracting the residuals from this model. This approach was necessary because ambient temperature increased throughout the period of sampling, and using residuals instead of raw temperature measurements allowed us to account for time of sampling on a particular day of sampling. Residuals from these models were used in all downstream analyses.

To perform subsequent analyses, we merged leaf morphology and area measurements from Migicovsky et al. (In Press) with residuals from the temperature model.

First, we performed a type 2 anova using the car R package v.3.0-11 (Fox and Weisberg, 2019). We used the following model (Equation 2), in which each of the principal component (PC) values are morphometric PCs calculated using the landmark data:

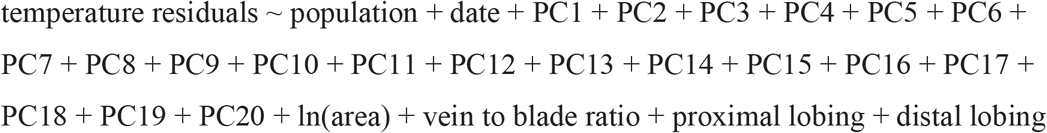

The first 20 morphometric PCs were used because culumultatively they explain 99.7% of the variance in leaf shape. The percent variation was calculated for all terms by calculating the Sum of Squares for a particular term, divided by the Total Sum of Squares, then multiplied by 100. The results for significant terms (p < .05) were plotted.

Since the highest amount of variance was explained by ln(area), scatterplots showing the relationship between ln(area) and the temperature residuals were plotted. In addition, to determine the correlation between these two measurements, a repeated measures correlation coefficient (r_rm_) was calculated. r_rm_ was used because correlation assumes independence of measures, but in this study we have four days of sampling of the same vines. The r_rm_ calculation accounts for this non-independence and was performed using the rmcorr R package version 0.4.5 (Bakdash and Marusich, 2021). Lastly, we used the weather data to calculate the average temperature, minimum temperature, and maximum temperature for each day of sampling.

## Results

In this study, we build on the findings of Migicovsky et al. (In Press) to explore the consequences of leaf shape variation on vine canopy temperature across 388 unique accessions resulting from five biparental crosses. Each biparental cross had one parent with compound leaves and a different second parent with non-compound leaves, and so these accessions vary primarily in the extent of lobing (Figure 1) but also differ in leaf area.

The canopy temperature of grapevines is both a critical indicator of water availability as well as having the potential to influence developmental timing/phenology. By recording both the time of sampling and vine canopy temperature, we were able to determine which vines were cooler or hotter than expected, and link this information with leaf morphology and size measurements.

The first objective of this study was to account for the time of sampling on canopy temperature, as measured using an infrared thermometer. To do this, we calculated a linear model for temperature ∼ date of sampling, and determined the value of each vine for a particular date, based on the residuals from that model (Figure 2). The slope of the line differed between dates, indicating that both date and time of sampling influenced canopy temperature.

**Figure 2.**
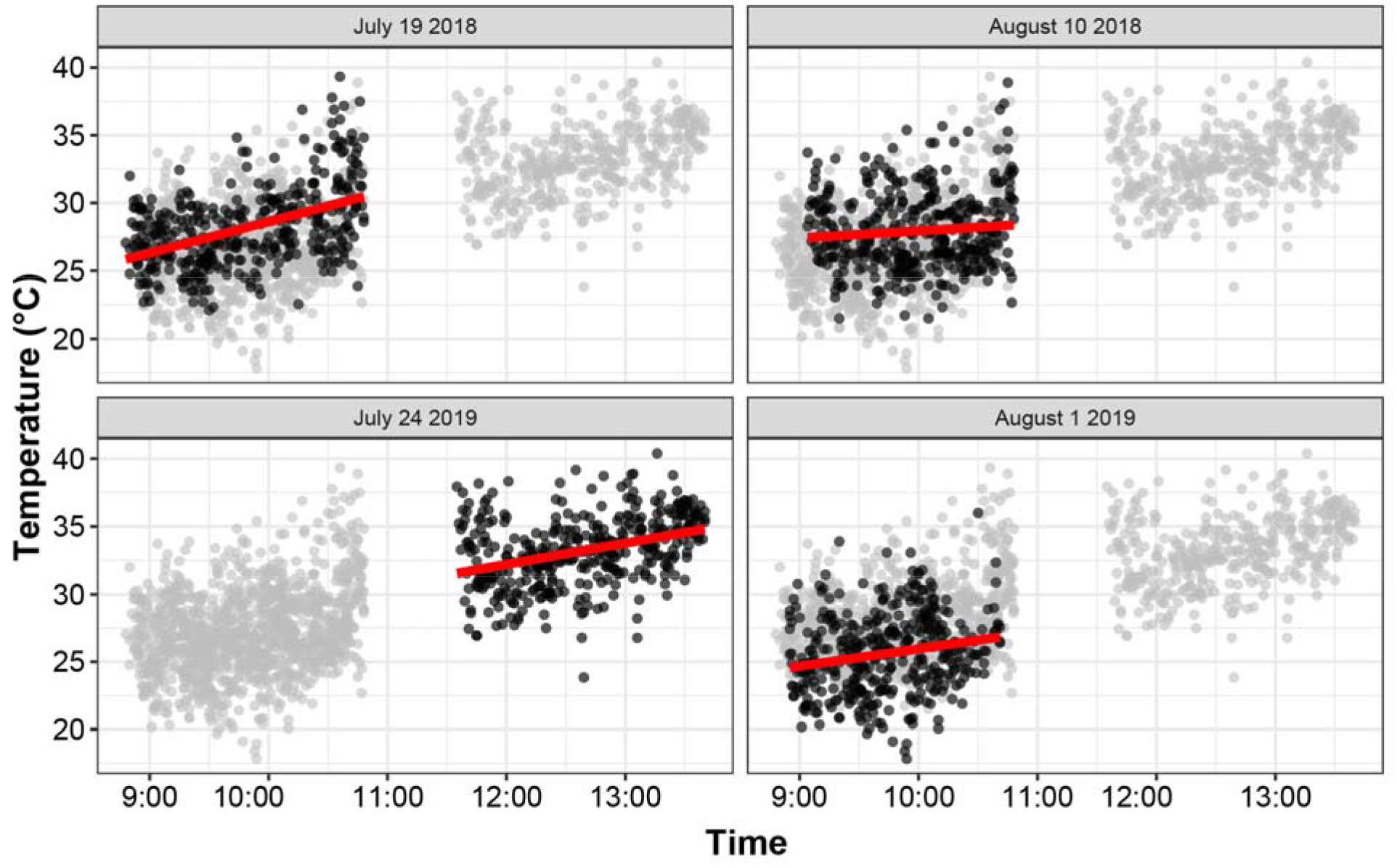
Scatterplots modeling canopy temperature vs time of sampling for each of the four dates measurements were taken. Each dot represents the temperature of a particular vine at a particular sampling day and time (n = 388). For each date, the measurements taken on that date are black, while the measurements from the remaining three dates are plotted in gray. The linear model for a particular date is shown using a red line.

After accounting for the time of sampling, we estimated whether leaf shape and size significantly influenced variation in temperature residuals. Temperature residuals were used because they allowed us to estimate whether a vine was cooler or hotter than expected, given the time of sampling. We performed a type 2 anova which accounted for morphometric PCs 1 to 20 (which cumulatively explain 99.7% of the variance in leaf shape), as well as leaf area on a natural logarithm scale, the vein to blade ratio, proximal lobing, distal lobing, which of the five populations the vines originated from, and the date of sampling. Across these factors, 4 were significant, including 3 morphometric PCs: PC14, PC18, and PC8, as well as leaf area (Figure 3). In all cases, less than 2.5% of the variance was explained by a given factor, with 0.5% or less explained for the morphometric PCs. Distal lobing, the primary source of variation in shape in the populations, was not significant (Migicovsky et al., In Press). In comparison, leaf area explained 2.13% of the variance in temperature residuals, which was the highest amount of any significant factors. Overall, these results indicate that leaf size, not leaf shape, has a stronger influence on variation in vine temperature.

**Figure 3.**
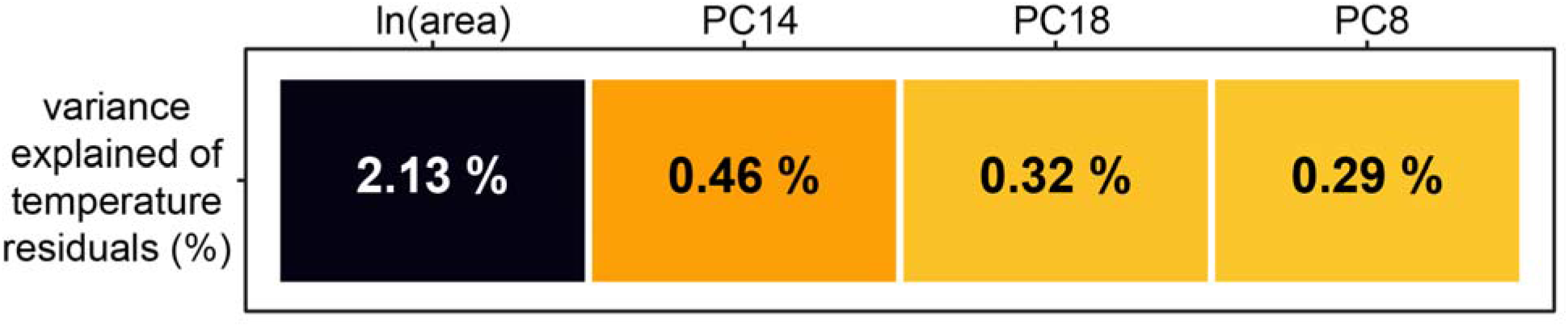
Percent variance explained by factors of interest estimated using a linear model (Equation 2) and type 2 anova. Only factors which explained a significant amount of variance (p <0.05) are included, with the color and text indicating the percent variance explained. Significant factors are sorted left to right from most variance explained to least variance explained.

Once we established a relationship between leaf area and canopy temperature, we used the repeated measures correlation coefficient to account for the non-independence of the four days of sampling and examined how the residuals from the temperature ∼ time model change in response to leaf area (Figure 4). We found that leaf area and the residuals were significantly negatively correlated (r = -0.178, p = 1.52 × 10^−12^). This negative correlation indicates that vines with larger leaves were cooler than expected, given the time of sampling.

**Figure 4.**
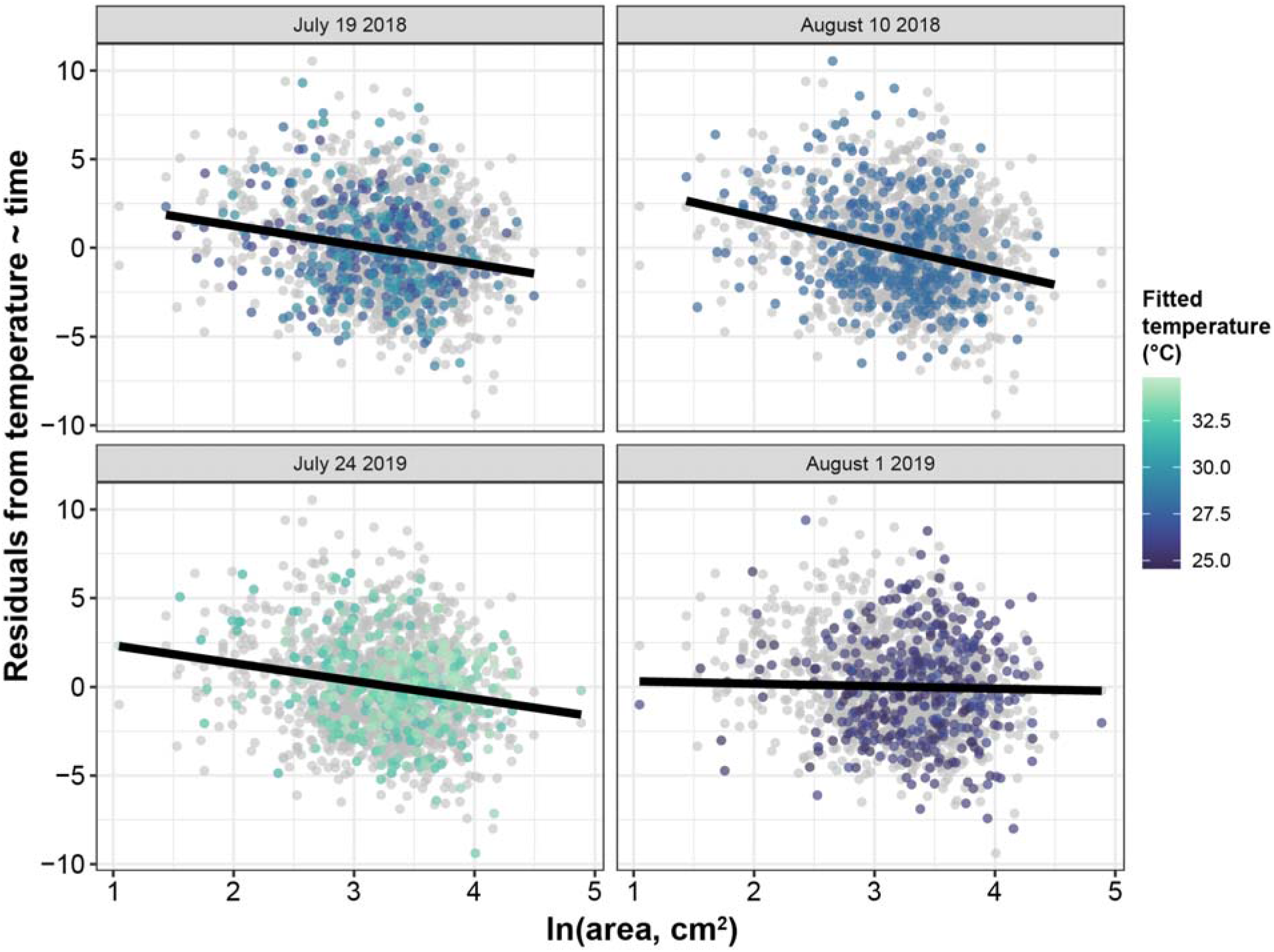
Scatterplot modeling the relationship between ln(area) and residuals from the linear model for temperature ∼ time. Each dot represents the measurement of a particular vine at a particular sampling day and time (n = 388). For each date, the measurements taken on that date are shown in the colour of the fitted temperature value, as shown in Figure 2, while the measurements from the remaining three dates are plotted in gray. The linear model for a particular date is shown using a black line, but the overall correlation was calculated using r_rm_ to account for the non-independence of the four days of sampling.

While the overall correlation is significant, it is clear that the relationship is strongest on the first three sampling days and not present on the final date of August 1 2019. When examining the weather data for these dates, the average temperature on August 1 2019 was cooler than the other 3 dates (24.6 °C in comparison to 26.9 to 28.1 °C) with the coolest max temperature values of 34.5 °C in comparison to 37.3 to 39.1 °C. A visual examination of the fitted temperature values which were adjusted based on time of sampling confirms that on August 1 2019 the canopy temperatures were the lowest (Figure 4).

## Discussion

On a global scale, smaller leaves are generally found at drier sites in warm regions in comparison to large-leaved species which are found in wet and hot environments. In wet and cold environments, species with smaller leaves predominate (Wright et al., 2017). The difference for leaf sizes based on access to water is due to the thicker boundary layer that large leaves have, which makes them more reliant on transpirational water loss for cooling (Gates, 1968; Wright et al., 2017). Although the climate in Madera County, California is dry and hot, vines were fully irrigated and thus water was not a limiting factor. The cooler temperature of the canopy for vines with larger leaves may indicate that transpirational cooling was occurring at a higher rate, reducing the risk of high temperatures more efficiently in comparison to vines with smaller leaves. This relationship seems particularly probable when considered in the context of the ambient temperatures on days of sampling: on the coolest day, August 1 2019, the temperature was on average over 2 °C cooler than any of the other days, with a maximum temperature reached that was 2.8 to 4.6 °C less than the three other dates. This was the same day of sampling when we did not observe a relationship between leaf size and temperature residuals, indicating this trend is strongest on hotter days, when the transpirational cooling benefit provided by large leaves may be greater. In comparison, larger leaves may not provide the same benefit on cooler days when the vines are under less stress from temperature.

In one study of an Australian heat wave, some vines were covered with a protective layer to reduce heating while others were exposed. In exposed vines, transpiration increased by nearly three times while photosynthesis was reduced by 35%, delaying ripening and causing a reduction in berry quality (Greer and Weedon, 2013). These results clearly indicate the negative impact that excessive increases in canopy temperature can have on vine health and berry quality. The ability to maintain a cooler canopy temperature on hot days is desirable for grape growers, and our preliminary findings indicate that this may be possible with larger leaves.

While we measured both leaf size and shape in our study, we did not measure canopy architecture, photosynthesis, or water use efficiency, as the heterogeneity of individual seedling canopies is very high, and this substantially complicates these observations on unreplicated seedlings. Therefore, it is possible that while overall reduction in vine canopy temperature occurs with larger leaves, this may not be due to leaf size, but rather, for example, differences in the openness of the canopy. Future work making use of thermal remote sensing imaging (Still et al., 2021) would be particularly useful in order to estimate the canopy temperature across numerous vines at the same time, reducing the effect of timing and human error on the results. Indeed, thermal imaging partnered with the CWSI could ultimately facilitate precision viticulture by assessing water stress and the need for irrigation, and work in this area is ongoing (Tanda and Chiarabini, 2019). There are also numerous areas for further work based on this preliminary study, including research which captures both individual leaf size as well as total leaf area of vine.

## Conclusions

This study builds on our previous work which determined that more highly lobed leaves compensated for what would otherwise result in a reduction in leaf area by having longer veins and a higher vein to blade ratio (Migicovsky et al., In Press). In this study, we determined that vines with larger leaves had cooler canopies than anticipated. Taking these findings together indicates that it should be possible to select for large, highly lobed leaves with the corresponding benefits in reducing canopy temperature and improving photosynthetic capacity, while still allowing light to permeate the canopy. Given the established link between water status and canopy temperature, this study lays the groundwork for future studies examining the effect of grapevine leaf size and shape. Ultimately, if grape breeders could harness variation in the size of grapevine leaves for reduction in canopy temperature, this could serve as a valuable target for future cultivar improvement.

## Acknowledgments

The research conducted for this study was supported by the National Science Foundation (NSF) Plant Genome Research Program 1546869. JFS was supported by an NSF Graduate Research Fellowship under Grant No. 1758713, and Saint Louis University. This research is also funded through the USDA National Institute of Food and Agriculture and Michigan State University AgBioResearch.

We acknowledge all of the individuals involved in maintaining the vineyard evaluated for this study. We acknowledge Leah Brand (Missouri State University), Julie Curless (Missouri State University), Dalton Gilig (University of Missouri), and Ilona Natsch (Saint Louis University) for assistance in sampling and landmarking of the leaves. We would also like to acknowledge Laszlo Kovacs (Missouri State University) for student supervisory support.

## Data Availability

All data and code used in this study can be found on GitHub (https://github.com/zoemigicovsky/grape_leaf_temp). All original scans used in this study are available from Dryad (Migicovsky et al., 2022)

## Author Contributions

PC generated the seedlings and supervised the maintenance of the vineyard. ZM, JFS, PC, and DHC conceived of the initial idea for this study. ZM coordinated the research. ZM, JFS, ZH, LLK, AL, MM, and KW sampled the leaves for this study. AF, MK, AJM, PC, and DHC acquired the funding for this study and provided supervisory support. ZM performed the data analysis with input from DHC. ZM wrote the first draft of the manuscript, which all authors read, commented on, and edited.

